# Evidence of antibodies against SARS-CoV-2 in wild mustelids from Brittany (France)

**DOI:** 10.1101/2022.01.20.477038

**Authors:** Bernard Davoust, Patrick Guérin, Nicolas Orain, Camille Fligny, Fabien Flirden, Florence Fenollar, Oleg Mediannikov, Sophie Edouard

## Abstract

In the French region of Brittany, mainly in the department of the Côtes d’Armor, during the first semester of 2021, seropositivity for SARS-CoV-2 was detected in five wild mustelids out of 32 animals tested. Anti-SARS-CoV-2 IgG against at least four out of five recombinant viral proteins (S1 receptor binding domain, nucleocapsid, S1 subunit, S2 subunit and spike) were detected using automated western blot technique in three martens (*Martes martes*) and two badgers (*Meles meles*). An ELISA test also objectified seropositivities. Although the 171 qPCRs carried out on samples from the 33 mustelids were all negative, these preliminary results (observational study) nevertheless bear witness to infections of unknown origin. The epidemiological surveillance of Covid-19 in wildlife must continue, in particular with the tools of efficient serology.

## 1. Introduction

Human infection by a newly identified coronavirus, SARS-CoV-2, was reported in China, end of 2019 (Huang et al., 2020). This pathogenic coronavirus is responsible for the COVID-19 pandemic which, over two years, has caused, 288 million cases of infection and 5.45 million deaths [www.worldometers.info/coronavirus/]. Despite the health measures taken and the massive use of vaccines, SARS-CoV-2 continues to spread, particularly due to the appearance of new genetic variants. The precise origin of this virus has not yet been firmly established, but the fact that the coronavirus closest to SARS-CoV-2 (BatCoV RaTG13) has been identified in Chinese horseshoe bats (*Rhinolophus affinis*), enables us to hypothesize that it is a zoonotic pathogen (Zhou et al., 2020). An animal coronavirus has crossed the barrier passing from bats to humans via, possibly, another close-to-human animal acting as a vector or even a secondary reservoir. This passage from an animal coronavirus to humans has resulted in adaptation to the host through viral mutations. Today SARS-CoV-2 spreads primarily from person to person all over the world. Animals seems low implicated in the spread of Covid-19 (Maurin et al., 2021). First, animals can be infected by asymptomatic infected or sick people. This has been well described in domestic animals (cats, dogs, ferrets), animals raised for fur (minks) and zoo animals (felines, primates, etc.) (Maurin et al., 2021; Fenollar et al., 2021; Jemeršić et al., 2021; Pomorska-Mól et al., 2021; OIE, 2022). To date, cases of human infection with SARS-CoV-2 from an animal (reverse zoonosis) have proved exceptional and limited to the particular ecosystem of mink farms (Hammer et al., 2021; Oude Munnink et al., 2021). The latest epidemiological studies show that there are now wild animals naturally infected with SARS-CoV-2. These observations of a reservoir of pathogens, susceptible to mutations, and potentially responsible for transmission from wildlife to humans, are of great interest in the context of the prevention of Covid-19 (Delahay et al., 2021). Thus, white-tailed deer (*Odocoileus virginianus*) in the USA and then in Canada are reservoir hosts for SARS-CoV-2 (Palermo et al., 2021; Hale et al., 2021; Palmer et al., 2021). In addition, in wild American minks (*Neovison vison*) from Utah (USA) and Spain, a SARS-CoV-2 infection has been detected (Shriner et al., 2021; Aguiló-Gisbert et al., 2021). It is well known that mustelids are very receptive and sensitive to the point that the ferret has become the best model of Covid-19 for experimental infections (Alluwaimi et al., 2020; Boklund et al., 2021). In a farm in western France, minks were infected and then euthanised at the request of the health authority (Anses, 2021). In this context, an observational study, limited in time and space, was conducted to detect SARS-CoV-2 infection in wild mustelids collected in the French region of Brittany.

## 2. Materials and methods

### 2.1. Animals and samples

Following an agreement with the hunting federations of two French departments in Brittany, Morbihan and Côtes d’Armor, we were able to take samples from the corpses of 33 mustelids, just after their death. From April to June 2021, we sampled: 14 martens (*Martes martes*), 10 badgers (*Meles meles*), 4 American minks (*Neovison vison*), 3 polecats (*Mustela putorius*) and 2 beech martens (*Martes foina*). In seven cases, the animals died as a result of a road collision. In addition, 11 mustelids were shot dead in accordance with current hunting regulations and 15 others were trapped and shot down in application of article R 427-6 of the French Environment Code. For ethical reasons of biodiversity protection, strict limits were imposed on the number of animals studied. The total group of 33 mustelids studied was made up of 19 females and 14 males. There were 23 animals from the department of the Côtes d’Armor (No. 22) and 10 from the Morbihan (No. 56). The sites where the mustelids were found dead or shot were located by their latitude and longitude (Table 1).

**Table 1.**
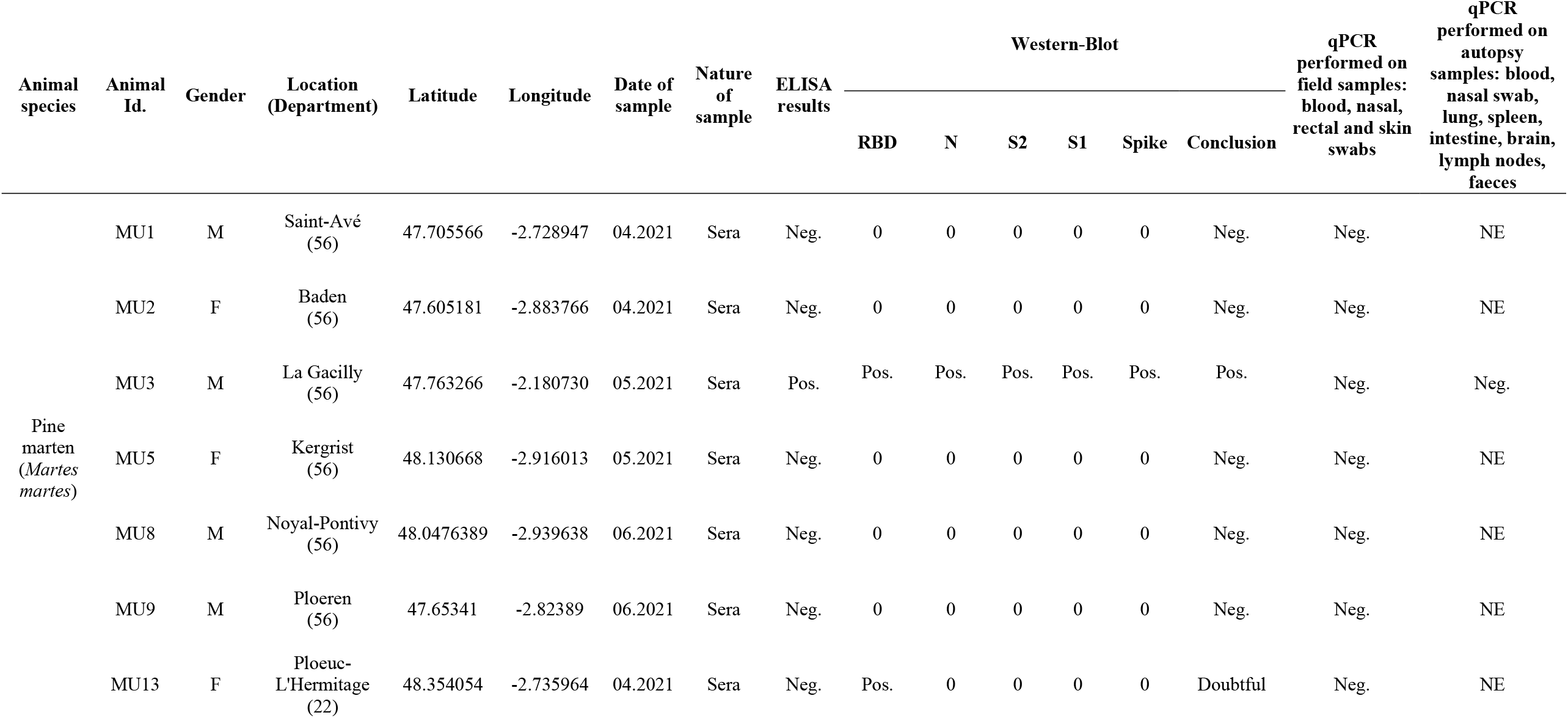
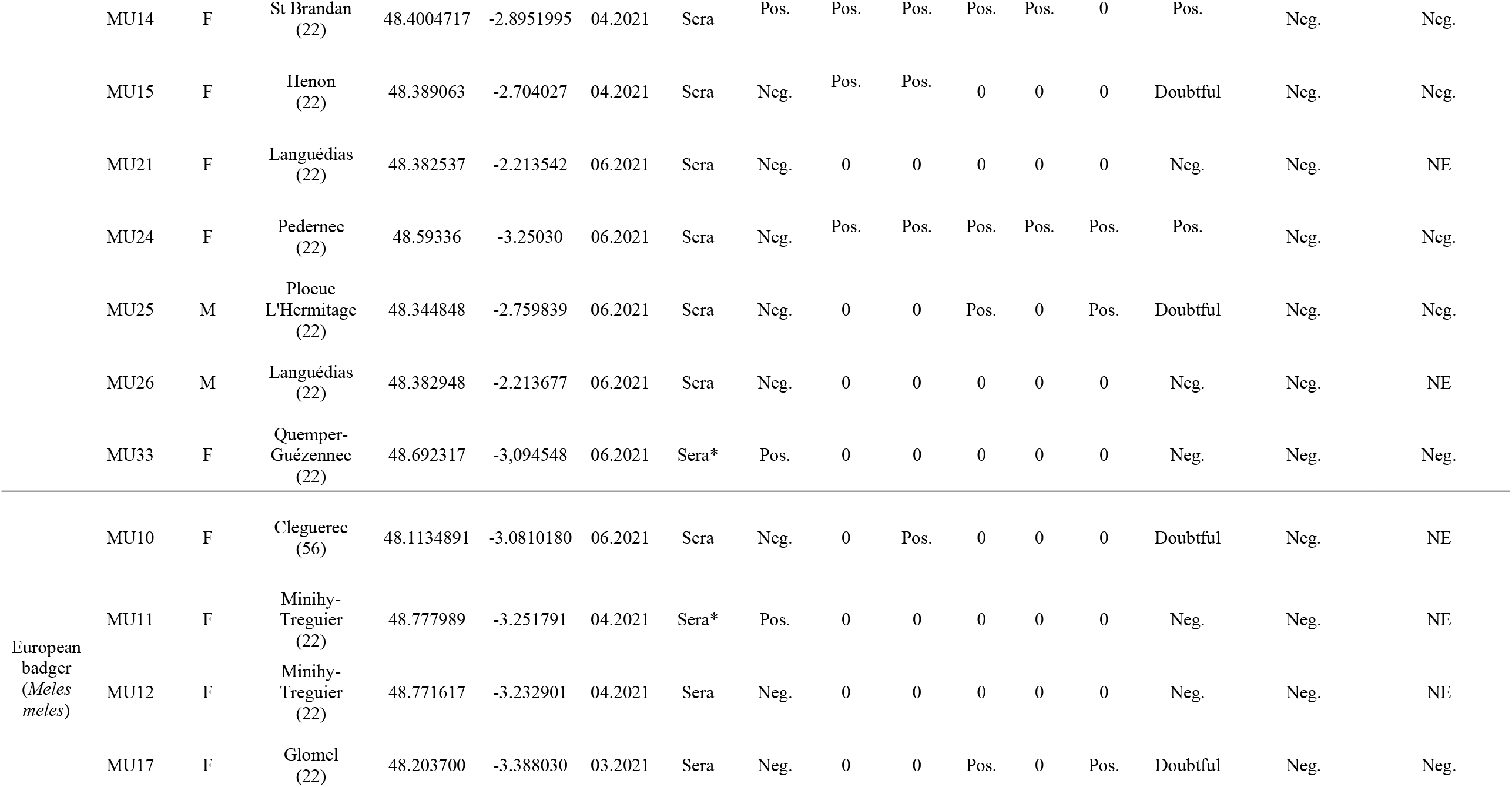
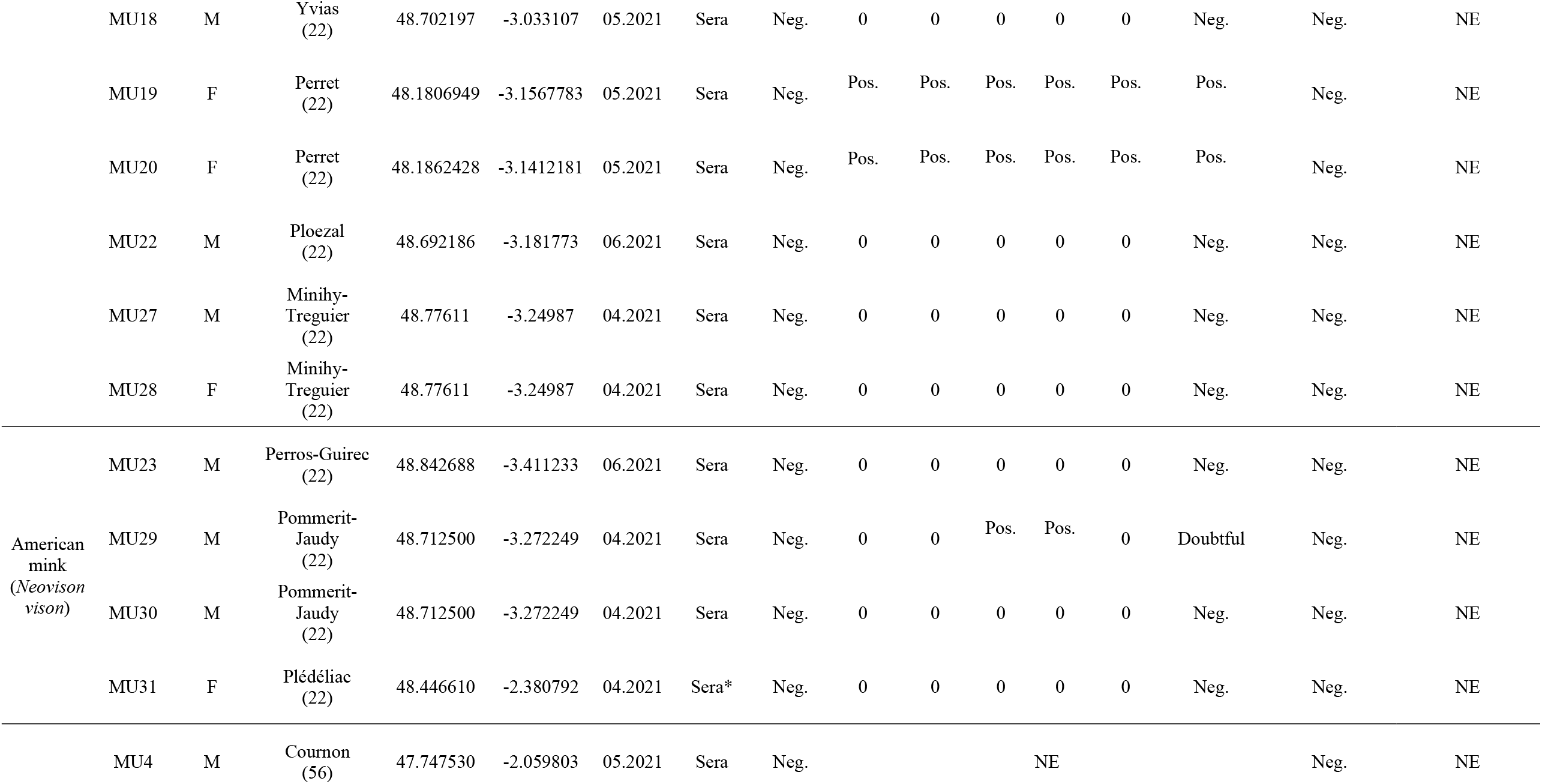
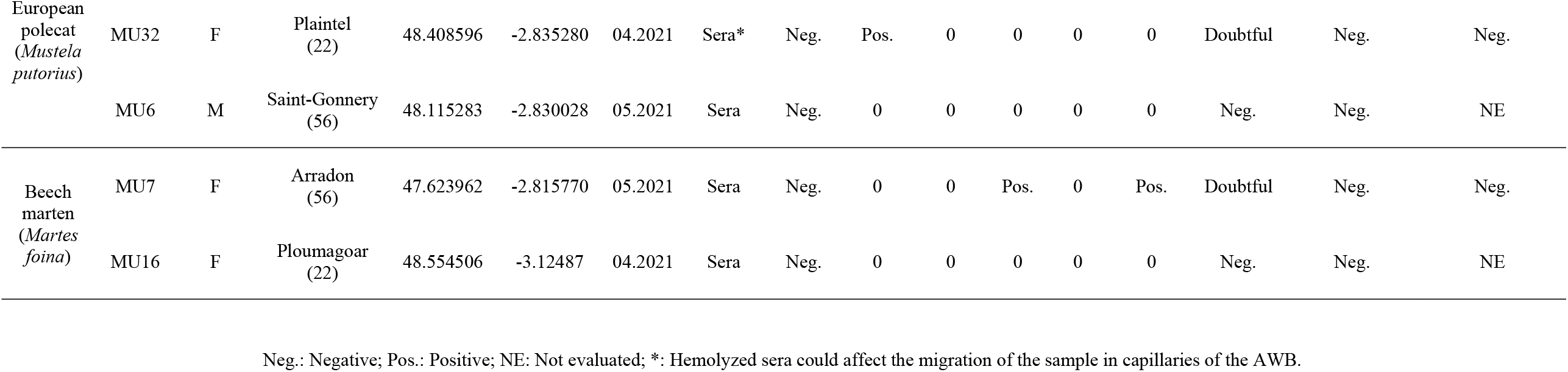
Results of SARS-CoV-2 ELISA, automated western immunoblotting and qPCR of 33 wild mustelids from Brittany (France).

In the field, we performed nasal, skin and rectal swabs as well as a blood sample from the heart (one tube of blood on EDTA and one dry tube with serum separator gel). These samples were transported at + 4 °C to the laboratory (IHU Méditerranée Infection, Marseille) in less than 48 hours. The corpses of the mustelids were then stored frozen at - 20 °C in Brittany. Subsequently, they were transported to the IHU for autopsy and tissue sampling: lung, spleen, intestine, brain, blood, faeces, nasal swab, and lymph nodes.

### 2.2. Serological detection

#### 2.3.1. ELISA assay

For ELISA, we used ID Screen^®^ SARS-CoV-2 Double Antigen Multi-species (Innovative Diagnostics, Grabels, France) following the manufacturer’s instructions. The test targets multispecies (i.e., minks, ferrets, cats, dogs, cattle, sheep, goats, horses and all other receptive species) total antibodies (IgG, IgM and IgA) directed against the major nucleocapsid protein of SARS-CoV-2. Plates were sensitised with a purified recombinant N antigen. Optical density (OD) was measured at 450 nm using Multiskan GO software (Thermo Scientific, Waltham, MA, USA). The test was validated when the optical density of positive control (OD_PC_) was ≥ 0.35 and a mean ratio of positive (OD_PC_) and negative (OD_NC_) control was higher than three. The optical density of each sample (OD_N_) was used to calculate the sample to positive (S/P) ratio (expressed as a %) where S/P= 100 * (OD_N_ - OD_NC_)/(OD_PC_ - OD_NC_). When the S/P score was lower than 50% by ELISA, samples were considered negative. They were considered as positive when it was higher than 60% and doubtful when 50< P/S score< 60%.

#### 2.3.2. SARS-CoV-2 antigen preparation and automated western immunoblotting (AWB) assay

The Jess^™^ Simple Western automated nano-immunoassay system (ProteinSimple, San Jose, CA, USA, a Bio-Techne Brand), a capillary-based size separation of proteins was used to evaluate the absolute serological response to five viral antigens from sera (Edouard et al., 2021). SARS-CoV-2 Multi-Antigen Serology Module® including S1 receptor binding domain (RBD) (48-kDa), nucleocapsid (58-kDa), S1 subunit (105-kDa), S2 subunit (71-kDa), and spike (170-kDa) recombinant proteins as antigens (ProteinSimple) and the 12-230-kDa Jess separation module (SM-W004) were used according to the manufacturer’s recommendations. Viral protein migration was performed through the separation matrix at 375 volts and were immobilised using photoactivated capture chemistry within the ProteinSimple proprietary system. Sera diluted at 1:2 were incubated for 60 minutes followed by a wash step and underwent 30 minutes incubation within an anti-ferret IgG conjugate (Abcam, Cambridge, UK) diluted to 1/200. The peroxide/luminol-S (ProteinSimple) was used for the chemiluminescent revelation. The Compass Simple Western software (version 6.0.0, ProteinSimple) was used for to capture the digital image of the capillary chemiluminescence and results analysis. A seropositive result with regard to SARS-CoV-2 is defined by an AWB Jess^™^ Covid-19 test showing reactivity against at least 4 out of the five recombinant proteins characteristic of SARS-CoV-2.

### 2.3. Biomolecular detection

Viral RNA extraction was performed on an EZ1 Advanced XL device using the EZ1 virus mini kit V2.0 according to the manufacturer’s recommendations (Qiagen, Courtaboeuf, France). The qPCR was run on a Lightcycler® 480 thermocycler (Roche diagnostics, Mannheim, Germany) using real time fluorescent RT-PCR kit for 2019-nCoV (BGI genomics, Hong Kong, China) targeting ORF1ab gene. Positive and negative (sterile water) controls were added in each qPCR runs and phage RNA internal control was added in each sample to validate RNA extraction and amplification (Amrane et al., 2020).

## 3. Results

All the results are presented in Table 1. Totally, positive ELISA was found in 4/33 (12.12% [IC 95%: 0.99; 23.26]) sera and positive AWB on 5/32 (15.6% [IC 95%: 3.04; 28.21]) sera. Five mustelids (four females and one male) were seropositive using AWB and showed high reactivity for RBD, nucleocapsid, S1 subunit, S2 subunit and/or spike (Figure 1). Positive mustelids were three martens (MU3, 14, 24) and two badgers (MU19 and 20). These two badgers were slaughtered at the same place, in Perret (Côte d’Armor), the same day in May 2021. Two martens were from the Côtes d’Armor and the third from Morbihan but from a town, La Gacilly, located on the southern edge of the Côtes d’Armor. In addition, two of these five mustelids (MU3 and 14) were also positive in ELISA. The sera of eight mustelids showed reactivity only against one and/or two proteins out of five viral proteins of AWB. The badger MU10 showed reactivity only against the nucleocapsid protein, MU32 and MU13 showed reactivity only against RBD. MU7, MU25 and MU17 showed reactivity against both spike and S2 protein, MU29 against S1 and S2 subunits and MU15 only against RBD and nucleocapsid. For the polecat (MU4), the AWB test was not performed due to insufficient serum. Furthermore, there were three false negative animals with the ELISA test (MU19, 20, 24) and two positive animals (MU11 and MU33) with ELISA for which AWB remained negative.

**Figure 1.**
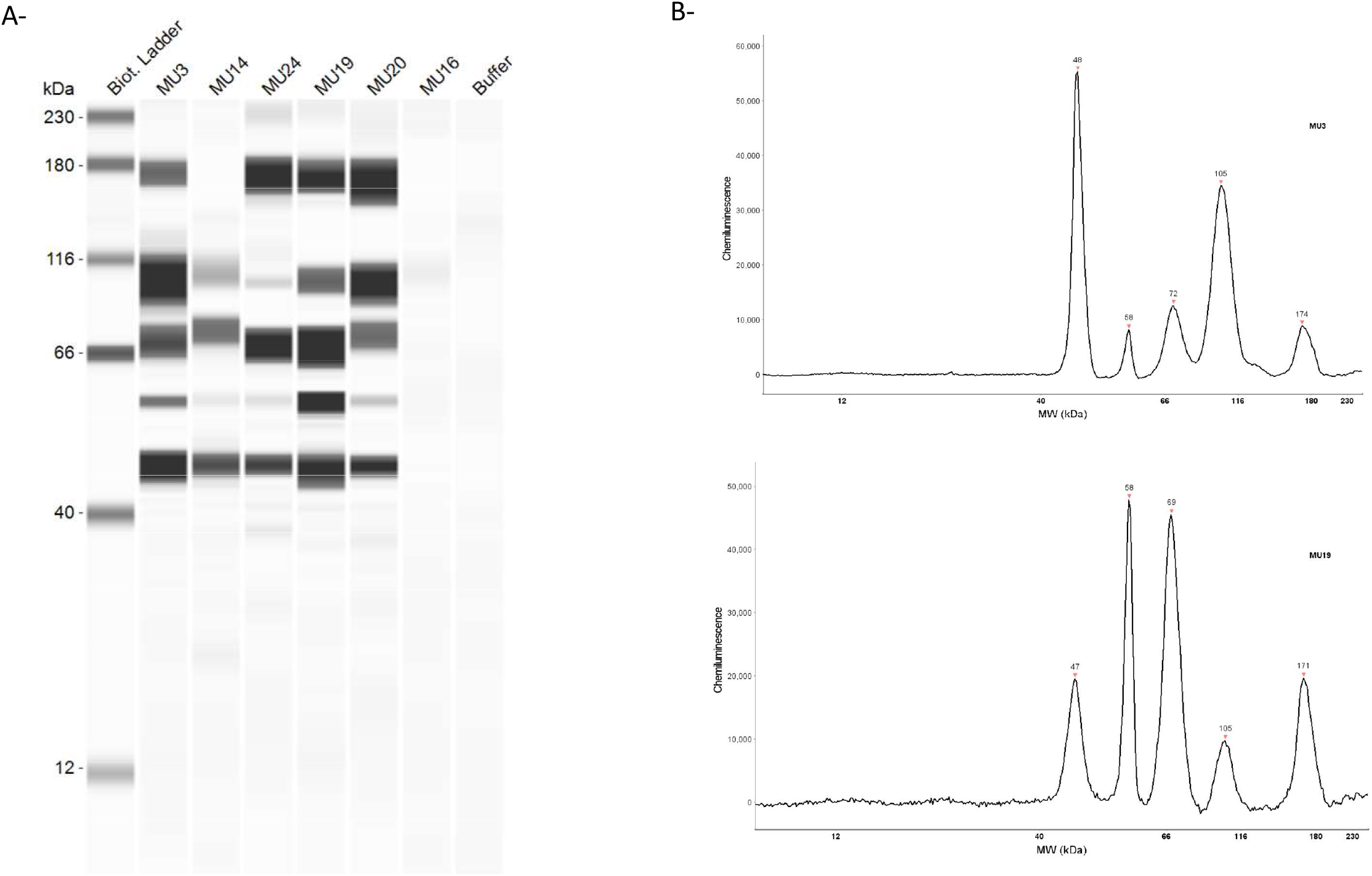
**(A)** Lane view of automated Western immunoblotting including the 5 positive wild mustelids (MU3, MU14, MU24, MU19, MU20) and one negative mustelids (MU16). The first lane represents the molecular mass marker in kDa and the last lane the negative control. **(B)** Chromatogram of chemiluminescence intensity detected by Jess^™^ Simple Western in the capillaries on positive pine marten (MU3) and on positive European badger (MU19). Bands and peaks were observed for S1 receptor binding domain (RBD) (48-kDa), nucleocapsid (58-kDa), S1 subunit (105-kDa), S2 subunit (71-kDa), and/or spike (170-kDa).

All the swabs (nasal, rectal and cutaneous) and the blood sample taken in the field on the 33 mustelids were negative in the specific SARS-CoV-2 qPCR test. Likewise, all the qPCRs carried out, a posteriori, on the samples taken from nine corpses kept frozen were negative.

## 4. Discussion

After discussing the reliability of our results, we will move on to their epidemiological significance. The potential impact on public health of transmission of SARS-CoV-2 to wild mustelids will be the subject of recommendations for active epidemiological surveillance.

Initially, serological screening was carried out with the ELISA test. Due to the positivity of several sera, additional investigations were implemented with the western blot technique which gives more precise results. The strong serological reactivity against four or five different antigens of the virus confirm specificity of antibodies to this coronavirus, indicating a humoral immune response linked to contact with the agent of the Covid-19 pandemic. All five AWB positive animals (MU3, 14, 19, 20, 24) were definitely infected with SARS-CoV-2. For eight mustelids, the AWBs were doubtful and were not considered positive for SARS-CoV-2. The AWB profiles showed reactivity against only one or two proteins suggesting cross reactivity with another coronavirus than SARS-CoV-2 or an uncompleted serological response (Lv et al., 2020; Li and Li, 2021). It is known that minks can be infected with an *Alphacoronavirus*, which is not zoonotic (Stout et al., 2021). Moreover, one ferret enteric coronavirus (FRECV) is similar to feline coronaviruses (Haake et al., 2020). The ELISA kit uses a truncated nucleocapsid protein in order to limit cross reactions with other coronaviruses (Spada et al., 2021). The diagnostic specificity of this test based on double antigens is > 99%, in dogs (Laidoudi et al., 2021). This ELISA test is useful to investigate SARS-CoV-2 antibodies in minks (Chaintoutis et al., 2021). Dissonant results were found between ELISA and AWB on five mustelids. Serological discrepant results were described among the different approach and are mainly due to antigen choice and sensitivity/specificity of the technique (Van Elslande et al., 2020). In the case of the badgers MU11 and MU33, the serum was too hemolyzed to draw reliable conclusions of AWB. Blood was drawn from the hearts of all corpses immediately after slaughter or within an unknown time frame after death for the seven animals found dead on the road. This explains why many sera were hemolyzed which made the AWB sometimes difficult to perform (capillaries blocked by red blood cells). Regarding our ELISA test, based on a double antigen, three negative results were obtained for the AWB positives (MU19, 20, 24). The AWB is a more sensitive technique than ELISA because it has a broad detection spectrum (targeting five SARS-CoV-2 proteins). Furthermore, the ELISA test detects total antibodies (IgG, IgM, IgA) against the virus core protein, whereas AWB detects only IgG. Recently, another multispecies ELISA based on the detection of antibodies against the receptor-binding domain (RBD) of SARS-CoV-2 has been developed with a specificity of 100% and sensibility of 98.3% (Wernike et al., 2020). In addition, the luciferase immunoprecipitation systems (LIPS), employing the spike, gave good results in experimentally infected ferrets (Berguido et al., 2021).

We did not want to conduct a seroprevalence survey according to representativeness criteria usually used in epidemiological surveys. We chose to do an observational study of a deliberately small number of wild mustelids, carried out without a reasoned sampling plan. Opportunistically, we benefited from the collection of corpses by hunters acting according to the standards in force in only two French departments of Brittany and for a limited time of four months. Ours is therefore a case study, not representative of the epidemiological reality concerning each of the five species of mustelids examined, in Brittany and even less throughout France. The seroprevalence (15.6% - 5/32) of SARS-CoV-2 found in wild mustelids should therefore not be overinterpreted. It should be considered as a warning indicator so that epidemiological field surveys can be carried out without delay according to the scientific methodology in force.

To place our observations in the context of two Breton departments during the first half of 2021, it will be essential to consult the official epidemiological database relating to the Covid-19 pandemic (ARS Bretagne, 2021). On March 30, the morbidity rate was respectively, 192 and 186 per 100,000 inhabitants, in the Côtes d’Armor and Morbihan (Santé publique France, 2022). In Brittany, at the outset of 2021, the B.1.160 variant was still in the majority. Then on February 15, the prevalence of the British variant Alpha (B.1.1.7) reached 54.4%. It rose to 98.8% at the end of May (Santé publique France, 2022).

The mustelids in our study are common in France (“Least Concern” status of the International union for the protection of nature) and considered as likely to cause damage in the two departments. Badgers are slaughtered by administrative order of the prefects. In France, the American mink population is not indigenous. It is due to escapes from mink farms for fur, especially in Brittany, in the 1960s (Léger et al., 2018). Mustelids adapt to a varied diet depending on the seasons (small mammals, worms, insects, fruits, etc.). Their way of life is discreet with preferentially nocturnal outings from their burrows. They live in fields and forest areas and approach human dwellings only occasionally. Pine martens live mostly in woods and are generally solitary. They move along the ground with nosy behavior. They frequent places of human passage such as woodpiles and forest roads in search of prey (Schwanz, 2000). The transmission of SARS-CoV-2 to wild mustelids may have occurred, initially, through indirect contact with an infected human through environmental contamination (wastewater? household waste? aerosols?). All the mustelids studied lived in anthropized and non-isolated rural areas. From one or more index cases, transmission spread directly between mustelids. This is certain for Perret’s two badgers (MU19 and 20). Badgers have more social contact than martens (Wang, 2011). Another hypothesis regarding the origin of the infection can be made but has not been demonstrated: it is known that in November 2020, a mink farm in Eure-et-Loir was widely infected with SARS-CoV-2, which led to the slaughter of all the animals (Anses, 2021). It seems possible that an infected mink escaped from the farm and subsequently infected wild mustelids, via an epidemiological chain of transmission. The epizootic could have spread in a few weeks, in particular, as far as the Côtes d’Armor, located some 300 km away. This distance renders this hypothesis rather improbable. Nevertheless, escapes of this type have been strongly suspected for the outbreaks in Utah (USA) and Spain (Shriner et al., 2021; Aguiló-Gisbert et al., 2021).

Viral circulation between mustelids is rapid. Indeed, they are the most receptive animals to SARS-CoV-2 (Shuai et al., 2020). Like humans, they have the angiotensin-converting enzyme 2 (ACE2) receptor on the cells of the respiratory tract, which facilitates viral penetration (via the spike protein) and infection (Covid-19) (Lean et al., 2021). Infection with SARS-CoV-2 of ferrets (*Mustela putorius furo*), laboratory animals, shows that they remain carriers of the virus for 14 days while the specific antibodies persist for several months (Monchatre-Leroy et al., 2021). In addition, in infected farms, minks were generally asymptomatic, some of them present with cough and fever. Excess mortality can be observed (Boklund et al., 2021; Pomorska-Mól et al. 2021).

The seropositivities that we have highlighted are proof that wild mustelids are good epidemiological sentinels for Covid-19. The question of their role as a reservoir for SARS-CoV-2 must be asked even if the five seropositive animals were no longer carriers of the virus. The problematic with mustelid coronaviruses is their mutagenic power, which produces potentially zoonotic viral variants. This has been well documented in a mink farm in Denmark (Oude Munnink et al., 2021; Hammer et al., 2021). Twelve people in contact with mink carrying SARS-CoV-2 were infected with an entirely new emerging variant (cluster 5) (Lassaunière et al., 2021). This FVI-spike variant virus corresponds to a combination of four mutations (69-70-deltaHV, 453F, 692V and 1229I) (Bayarri-Olmos et al., 2021; Lassaunière et al., 2021). This outbreak was quickly brought under control but remains a model for understanding the health risk associated with mustelids, reservoirs of coronaviruses transmissible to humans. In addition, very recently, it was shown by the study of the genome and the mutations that the Omicron variant (B.1.1.529) of SARS-CoV-2 could originate from a human virus passed through the mouse in which it would have mutated and then it would have again infected man (Wei et al., 2021).

To protect public health, reinforced epidemiological surveillance and biosecurity measures have been taken, at the request of the health authorities, in the three mink farms remaining in France (Anses, 2021). In addition, it is likely that mink farming will be banned in France in the future. However, our study shows that the pandemic virus circulates in wildlife in mustelids of two species (marten and badgers). It is therefore important to extend our one-off investigation to an active epidemiological surveillance of mustelids (injured or slaughtered) from corpses collected by the departmental hunting federations. Implementing this behoves the ministries responsible for wildlife and animal diseases.

The infection of wildlife with SARS-CoV-2 has been increasingly studied in the United States since the discovery of infected white-tailed deer (*Odocoileus virginianus*) (Palermo et al., 2021; Hale et al., 2021). Antibodies were detected in 152 samples (40%) from 2021 using a surrogate virus neutralization test (Chandler et al., 2021). The role of these deer in the evolution of the pandemic is not known. From the outset of the pandemic, domestic animals (dogs and cats) were found to be carriers of SARS-CoV-2 and have been the subject of numerous studies and case reports (OIE, 2022). With the identification of cases in wildlife, the OIE has proposed specific recommendations (OIE, 2020). Their application will make it possible in the future to better understand the epidemiological situation in several countries.

## 5. Conclusion

This pilot study, limited in time and space, can only provide ad hoc information. However, it is decisive from the point of view of the circulation of a zoonotic virus which, until now in France, had only been detected in humans, dogs, cats and farmed mink. As Brittany is not a particular ecosystem for mustelids, it is necessary to step up research through epidemiological surveys, in particular with the tools of efficient serology, in other regions and countries. The demonstration of real seroprevalences by species of mustelids will have to be based on very reliable serological methods such as the Jess^™^ automatic western blot that we used coupled with an ELISA. From the origin of the virus in China at the end of 2019, until today with the circulation of SARS-CoV-2 in the wildlife of other continents, animals have been observed to play a role that should not be neglected in the pandemic of Covid-19. In a *One Heath* approach, it is therefore necessary to intensify cooperation between physicians, veterinarians and all professionals of domestic animals and wildlife.

## Acknowledgements

The authors are grateful to the staff of OpenHealth Company and the Côtes d’Armor and Morbihan hunter federations who made the collection of samples possible. The authors would like to thank Innovative Diagnostics which provide them the ELISA ID Screen® SARS-CoV-2 kit. We are grateful also to Raphaël Tola, Laurence Thomas, Naomie Canard and Marie-Charlotte Mati for their technical help.

